# Comprehensive mapping of HIV-1 escape from a broadly neutralizing antibody

**DOI:** 10.1101/117705

**Authors:** Adam S. Dingens, Hugh K. Haddox, Julie Overbaugh, Jesse D. Bloom

## Abstract

Precisely defining how viral mutations affect HIV’s sensitivity to antibodies is vital to the development and evaluation of vaccines and antibody immunotherapeutics. But despite great effort, a full map of escape mutants has not yet been delineated for even a single anti-HIV antibody. Here we describe a massively parallel experimental approach to quantify how all single amino-acid mutations to Envelope (Env) affect HIV’s sensitivity to a neutralizing antibody. We applied this approach to PGT151 and identified novel sites of escape in addition to those previously defined by structural and functional studies, such as glycans at sites 611 and 637, residue 647, and sites in the fusion peptide. Evaluating the effect of each amino acid at each site lends insight into the biochemical basis of escape throughout the epitope. Thus, comprehensive mapping of HIV antibody escape gives a quantitative, mutation-level view of the ways that Env can evade neutralization.

## Introduction

HIV’s rapid evolution enables it to outpace even the exceptional adaptive capacity of the humoral immune system. Although the immune system is usually outmatched in this evolutionary arms race, leading to viral persistence, infected individuals occasionally develop antibodies capable of neutralizing diverse viral strains. While these broadly neutralizing antibodies (bnAbs) do not control infection in the individual in whom they arise, their identification has motivated efforts in rational vaccine design and antibody immunotherapeutics. For example, epitope mapping of bnAbs has revealed conserved sites of vulnerability on HIV’s envelope glycoprotein (Env), and a leading vaccine strategy is to design immunogens that elicit an antibody response targeting these conserved sites (Wu and Kong, 2016). bnAbs are also being tested in both prophylactic and therapeutic settings. Numerous studies in animal models have shown proof-of-concept that passively infused bnAbs can protect against infection (Pegu et al., 2017) and therapeutically suppress viremia during infection (Margolis et al., 2017). Similar bnAb-based immunotherapies are being tested in humans, with some early studies showing a transient reduction in viral load or delay of viral rebound after treatment interruption in some individuals (Bar et al., 2016; Caskey et al., 2015, 2017; Lynch et al., 2015a; Scheid et al., 2016).

Despite the impressive breadth and potency of bnAbs *in vitro,* HIV can eventually evade them *in vivo.* Viral isolates from individuals who develop bnAbs are typically resistant to neutralization, and resistance arises when bnAbs are administered to infected animal models (Klein et al., 2012; Shingai et al., 2013) or humans (Bar et al., 2016; Caskey et al., 2015, 2017; Lynch et al., 2015a; Scheid et al., 2016; Trkola et al., 2005). It is therefore important to prospectively identify potential bnAb escape mutations in order to best predict the outcome of their therapeutic use. However, this can be challenging, in part because bnAbs often target complex conformational and glycosylated epitopes. To date, a complete set of HIV escape mutations has yet to be elucidated for any antibody.

The limited observational studies of viral escape from bnAbs to-date likely reveal only a fraction of the full repertoire of escape mutations. Structural studies provide atomic-level views of the antibody-antigen footprint, but fail to reveal which interactions are necessary for neutralization and which mutations disrupt these interactions. Indeed, it has long been appreciated that binding energetics are often concentrated at select sites in the protein-protein interface (Clackson and Wells, 1995; Cunningham and Wells, 1993), and mutations at Env residues that participate in crystal-structure-defined interactions do not always affect bnAb binding and neutralizing (Falkowska et al., 2012; Li et al., 2011). Because structures do not functionally define escape mutations, researchers often generate and interrogate single amino-acid mutants in binding or neutralization assays. This approach is so labor intensive that it has only been applied to a fraction of the sites in Env, and typically to only one or a few mutations – often to alanine – at these sites.

We have applied a new deep mutational scanning-based approach to comprehensively map all mutations to Env that enable HIV to escape from a broadly neutralizing antibody. This approach, mutational antigenic profiling, involves creating libraries of all single amino-acid mutants of Env in the context of replication-competent HIV (Haddox et al., 2016), selecting for mutations that promote antibody escape, and using deep sequencing to quantify the enrichment of each mutation. It is conceptually similar to a strategy recently used by Doud et al. (Doud et al., 2017)to completely map the escape of influenza A from anti-hemagglutinin antibodies. Our work comprehensively profiles HIV escape from bnAb neutralization in the context of actual virus, distinguishing it from previous deep mutational scanning experiments that have used yeast or cell-surface display to identify Env variants that bind germline-encoded bnAb progenitors (Jardine et al., 2016; Steichen et al., 2016). The map of escape mutations that we generate provides a complete view of the functional interface between Env and a bnAb, and suggests biochemical mechanisms of escape that are not readily apparent from the incomplete data provided by traditional mapping approaches.

## Results

### Overview of mutational antigenic profiling

To define all single amino-acid mutations to Env that increase resistance to antibody neutralization, we implemented the mutational antigenic profiling approach in Figure 1. This approach relies on creating libraries of full-length proviral clones encoding all possible mutations in *env,* and passaging this virus in T cells (Haddox et al., 2016). The result is a library of replication-competent HIV encoding all functionally tolerated mutations in Env. This mutant library is then incubated with and without antibody and used to infect target cells. After entry, the frequency of each mutation in the infected cells is quantified by deep sequencing (Fig 1A). The *differential selection* that the antibody exerts on a mutation is defined as the logarithm of that mutation’s enrichment in the antibody-selected condition relative to the control, and selection for enriched mutations is plotted in logoplots as shown in Figure 1B. The entire experiment was performed in biological triplicate beginning from independent generation of the proviral plasmid mutant libraries (Figure 1C).

**Figure 1.**
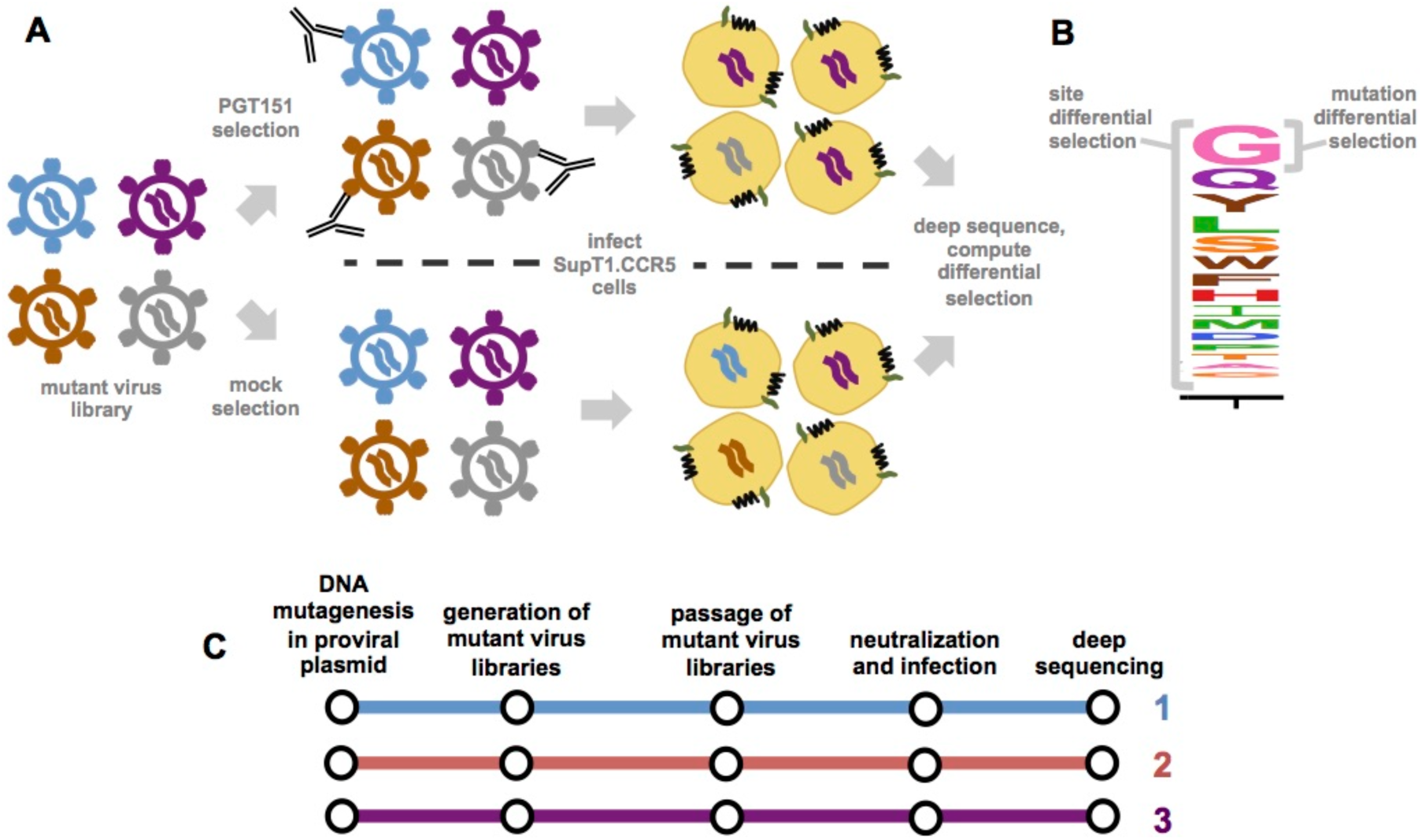
Schematic of mutational antigenic profiling. **(A)** Mutant virus libraries of HIV, which have been passaged in SupT1.CCR5 cells and thus should encode all single amino-acid mutants to Env compatible with viral replication, are incubated with and without an anti-HIV antibody, and then infected into SupT1.CCR5 cells. After viral entry and reverse transcription, cDNA is isolated and *env* is deep sequenced. The differential selection exerted by the antibody is quantified as the logarithm of the frequency of each mutation relative to wildtype in the antibody-selected condition compared to the control condition. **(B)** A logoplot visualizing the differential selection at a single site. The height of each letter is proportional to the differential selection for that amino acid mutation. The site differential selection is the sum of all mutation differential selection values at that site. **(C)** The entire mutational antigenic profiling process was completed in biological triplicate, starting from generation of three independent Env mutant libraries in the context of proviral plasmids.

### Generation and deep sequencing of mutant virus libraries

We applied mutational antigenic profiling to a virus relevant to HIV transmission by generating the mutant libraries in the context of an *env* (BF520.W14M.C2) from a subtype A virus isolated shortly after mother-to-child transmission in an infant who went on to rapidly develop a bnAb response (Goo et al., 2012; Simonich et al., 2016; Wu et al., 2006). We introduced all possible codon-level mutations to the gp120 surface unit and extracellular portion of gp41 (Env residues 32 to 703) in the context of a proviral plasmid. We did not mutagenize Env’s signal peptide or cytoplasmic tail, as these are not direct targets of neutralizing antibodies; further, they play a role in regulating Env cell surface expression (Chakrabarti et al., 1989; Li et al., 1994; Yuste et al., 2004). Sanger sequencing of individual proviral clones revealed that the number of mutations per clone followed a Poisson distribution with a mean of 1.1 mutations/clone, and the mutations were relatively uniformly distributed along the length of Env (Figure S1 and Figure S2).

We used deep sequencing to measure the frequency of each mutation in the mutant proviral plasmid, as well as in the mutant virus libraries with and without antibody selection. Because any given mutation is rare, we performed deep sequencing using a barcoded-subamplicon approach that tags individual molecules with unique molecular identifiers during the library preparation to increase accuracy (Doud and Bloom, 2016; Jabara et al., 2011; Wu et al., 2016). Deep sequencing of the mutant proviral plasmids showed that 95% of the 12,559 possible single amino-acid mutations (661 mutagenized residues × 19 mutant amino acids) were present in each of the triplicate BF520 plasmid mutant libraries, with 99% present in the three libraries combined. Figure S2 shows that the sequencing error rate is 1.5×10^-4^ mutations/codon (equivalent to 5×10^-5^ mutations/nucleotide) as assessed by sequencing unmutated wildtype proviral plasmid. This error rate is over 10-fold lower than the frequency of codon mutations in the plasmid mutant libraries.

We also quantified the rates of errors associated with viral replication by sequencing wildtype viruses passaged in parallel to each mutant virus library (described below). The rate of errors in these passaged wildtype viruses remained well below the rate of mutations in the virus mutant libraries, allowing us to reliably assess mutation frequencies (Figure S2). In the analyses below, we statistically correct for these errors associated with viral replication as described in the Methods.

### Comprehensive profiling of escape from antibody PGT151

We chose to profile escape from the bnAb PGT151, which targets a quaternary, cleavage-dependent epitope made up of glycans and protein residues in the gp120/gp41 interface and fusion peptide. This bnAb has been extensively studied, using both structural (Blattner et al., 2014; Lee et al., 2016) and functional approaches (Falkowska et al., 2014; Van Gils et al., 2016; Kong et al., 2016; Wibmer et al., 2017).

To generate mutant virus libraries, we transfected the BF520 mutant proviral plasmids into 293T cells and passaged the transfection supernatant in SupT1.CCR5 cells at a low MOI to establish a genotype-phenotype link and purge non-functional mutants. The resulting passaged mutant virus libraries, which we expect to carry virtually all Env amino-acid mutations compatible with viral replication, were used to infect cells in the presence and absence of PGT151. We then isolated and deep sequenced viral cDNA from infected cells to determine the frequency of each mutation, and computed the differential selection across Env.

Among the three biological replicates, the site differential selection was well correlated (R= 0.59-0.71; Figure 2A, Figure S3), indicating that the high-throughput experiments yielded reproducible results even when starting with fully independent mutant proviral plasmid libraries.

**Figure 2.**
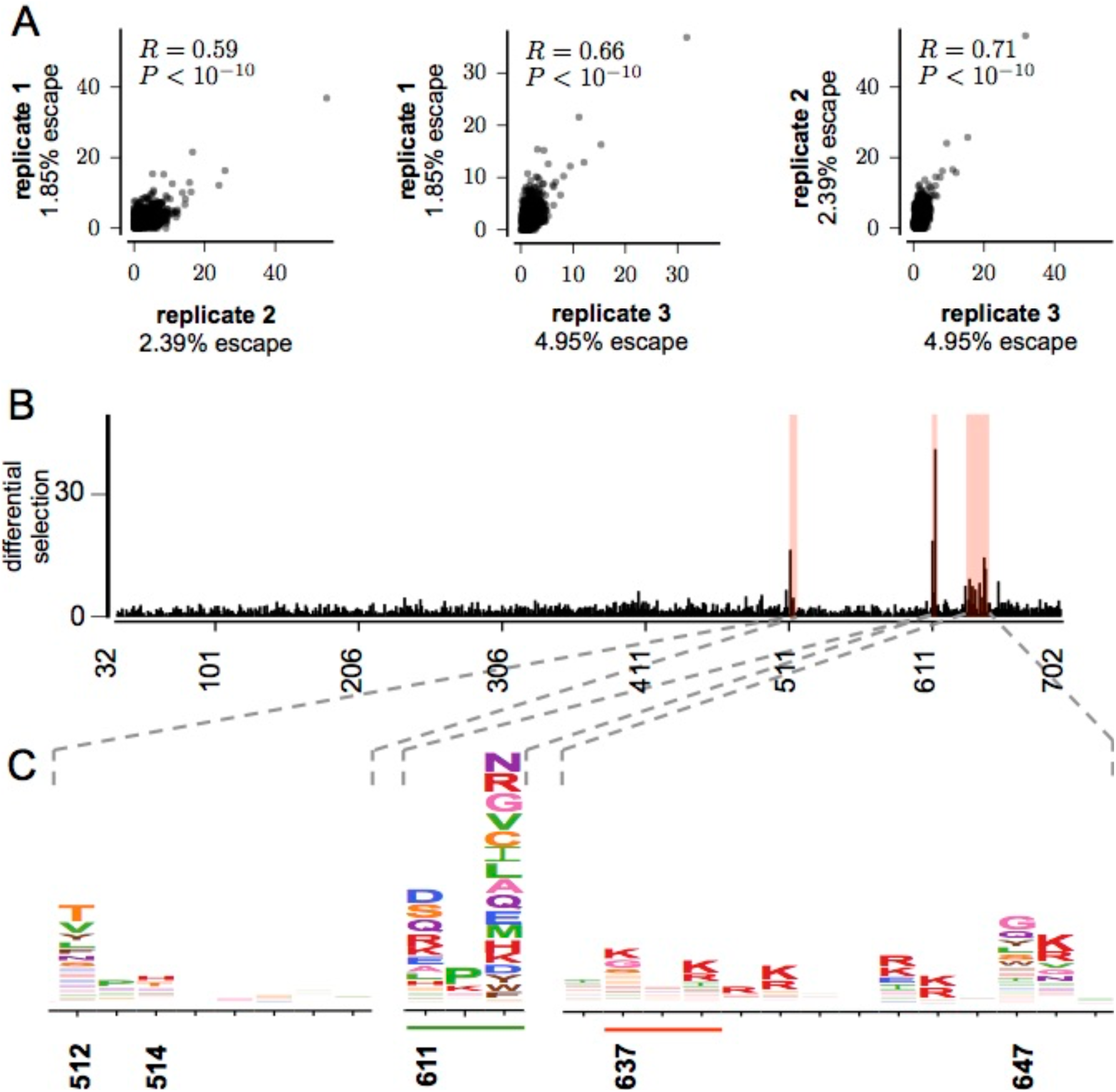
Reproducibility of mutational antigenic profiling. **(A)** Correlation of site differential selection across biological replicates. Each point indicates the selection at one of the 661 Env sites; plots also show the Pearson correlation and associated *P* value. Percent escape was calculated by using droplet-digital PCR to measure the number of viral genomes in the antibody-selected versus control sample. **(B)** The site differential selection, averaged across replicates, is plotted across the Env sequence. See Figure S3 for data from individual replicates. **(C)** Logoplots showing mutation differential selection in strongly selected regions. Mutations at numbered sites have previously been shown to reduce PGT151 neutralization sensitivity in other viral strains. From left to right: the fusion peptide, the 611 glycosylation motif (underlined in green), and the 637 glycan motif (underlined in red) and HR2 domain.

There was strong selection at sites in the known PGT151 epitope. Figure 2B plots the site differential selection across Env, and Figure 2C displays the escape profiles in regions of strong differential selection. These sites include residues originally mapped by mutagenesis and pseudovirus-neutralization assays: the dominantly targeted 611 glycan, the 637 glycan, and residue 647 (Figure 2B, 2C) (Falkowska et al., 2014). There was also strong differential selection at fusion-peptide sites 512 and 514, which have been mapped as part of the PGT151 epitope by structural (Lee et al., 2016) and functional (Van Gils et al., 2016; Kong et al., 2016; Wibmer et al., 2017) methods (Figure 2C). Therefore, our mutational antigenic profiling identified strong selection at epitope sites that have been identified by other approaches.

### Validation of mutational antigenic profiling at previously mapped sites

We next used TZM-bl neutralization assays of BF520 psuedoviruses with select mutations that were and were not differentially selected in our mutational antigenic profiling to further examine how well our comprehensive escape profiles predicted neutralization escape (Figure 3, S4). We first focused on sites that had been previously defined as conferring escape from PGT151, including mutations that disrupt the gp41 glycans at sites 611 and 637. Mutational antigenic profiling revealed strong selection for most mutations that disrupt the 611 glycosylation motif (N-X-S/T, where X can be any amino acid except proline) (Figure 3A). Both the 611 and 637 glycans make extensive contacts with the PGT151 Fab (Lee et al., 2016), and elimination of either can increase the IC_50_ in a number of strains (Falkowska et al., 2014). Of note, there was selection for proline at the central position of the 611 glycosylation motif, but not at the central position in the 637 glycosylation motif (Figure 3A). The 638P mutation was depleted in the mutant virus libraries passaged in the absence of PGT151 selection (data not shown), and we were unable to generate pseudoviruses bearing this mutation, suggesting that proline is not tolerated.

**Figure 3.**
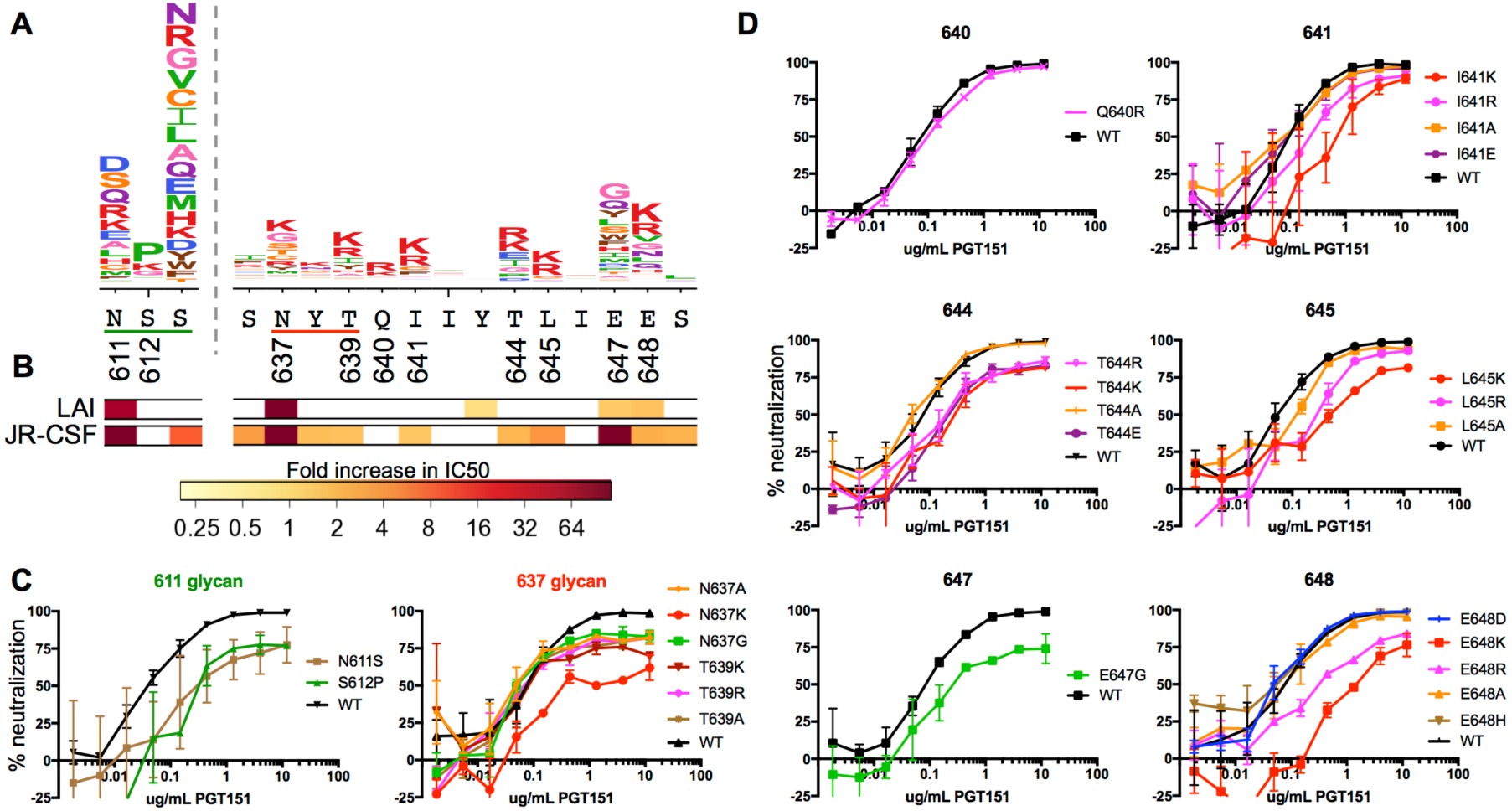
Analysis of the effect of individual amino acid changes identified by mutational antigenic profiling on PGT151 neutralization. **(A)** Logoplots showing differential selection. Sites where individual mutations were tested in neutralization assays are numbered, and the 611 and 637 glycosylation motifs are underlined in green and red. **(B)** Color bars below the logoplots indicate the results of neutralization assays performed on point mutants (primarily to alanine) of JR-CSF and LAI by Falkowska et al. 2014. The bar at each site is colored according to the fold change in IC_50_ relative to wildtype that a mutation imparted. If multiple amino acids were tested at a site, the site is colored according to the largest effect. **(C)** Results of TZM-bl pseudovirus neutralization assays on BF520 Env point mutants that alter the N-linked glycosylation motifs at sites 611 and 637. **(D)** Results of neutralization assays at individual residues outside of glycosylation motifs. In panels C and D, each plot shows a representative neutralization curve; see Figure S4 for regression curves from duplicate assays.

At site 637, some of the glycan knockout mutations are under stronger differential selection than others (Figure 3A). Consistent with the mutational antigenic profiling, the effect of these mutations on neutralization sensitivity in TZM-bl assays varied. According to the mutational antigenic profiling, the most strongly selected mutation at site 637 was to K (Figure 3A). In the neutralization assays, this mutation increased the IC_50_ by 3.7- fold. Disrupting targeted glycans can also alter neutralization sensitivity by decreasing the maximum percent neutralization; the 637 K variant also decreased the maximum percent neutralization to 59% (Figure 3C, 4A). All other tested mutations decreased the maximum percent neutralization of PGT151 to 74-81% (Figure 3C, 4A) but did not alter the IC_50_. These results agree with and expand on previous observations; mutations that disrupt the 637 glycosylation motif have been shown to decrease the maximum percent neutralization in JR-CSF, LAI, and JR-FL (Falkowska et al., 2014), and PGT151 often incompletely neutralizes pseudoviruses bearing many different Env variants (McCoy et al., 2015).

Prior work has also shown that mutations at site 647 reduce neutralization sensitivity in some strains (Figure 3B) (Falkowska et al., 2014). Mutational antigenic profiling agreed with these results, revealing strong differential selection for many mutations at site 647. We confirmed the most strongly selected mutation (E647G) mediated escape in a neutralization assay, with IC_50_ fold-change of 5.3 (Figure 3D).

Overall, these results show that mutations at previously identified sites that are also mapped by our high-throughput approach indeed have clear effects on neutralization sensitivity in the BF520 variant when tested individually.

### The comprehensive nature of mutational antigenic profiling allowed identification of new sites of escape

Our mutational antigenic profiling data showed strong selection at several sites where escape mutations have not previously been mapped. As noted above, both our work and prior studies identified escape mutations at site 647, where many different amino-acid mutations mediated escape (Figure 3A). In contrast, our mutational antigenic profiling also showed selection for mutations to positively charged amino acids at the neighboring site 648, where previous studies did not find effects of mutations to alanine or glycine (Figure 3A; (Falkowska et al., 2014)). Viruses engineered with E648K and E648R showed substantially reduced sensitivity to PGT151 (IC_50_ fold-change of 7.7 and 3.7, respectively), but mutations to a variety of other amino acids (D, A, and H) have no effect (less than 1.5-fold change in IC_50_) (Figure 3D).

At a number of additional positions in the heptad repeat 2 domain (HR2), including sites 641, 644, and 645, we also observed strong differential selection for predominantly positively charged mutations. Consistent with prior work in other strains (Falkowska et al., 2014), both our mutational antigenic profiling (Figure 3A) and pseudovirus neutralization assays (Figure 3D) showed no effect of alanine mutations at these sites. However, the positively charged escape mutations revealed by mutational antigenic profiling indeed reduced neutralization sensitivity when assayed in individual neutralization assays, with fold changes in IC_50_ ranging from 3.2-5.7 fold.

### Correlation between mutational antigenic profiling and traditional neutralization assays

At each validated site, the ranked order of effects of each mutation on neutralization sensitivity was well correlated between our high-throughput experiments and in individual TZM-bl assays (Figure 3, 4). Across all validated sites, we next asked if our differential selection measures were representative of the true neutralization phenotype. For the BF520 mutations that we tested, the enrichment measured by mutational antigenic profiling is well correlated with the fold-change in IC_50_ relative to wildtype from TZM-bl assays (R = 0.67, Figure 4). Remarkably, this is within the range of correlations observed between neutralization assays performed on the same sets of viruses by different laboratories (Todd et al., 2012). Further, the IC_50_ only partially summarizes the information in a neutralization curve, as mutations also altered the maximum percent neutralization (Figure 4A). Many of the mutations that exhibit the poorest correlation between fold change in IC_50_ and enrichment in mutational antigenic profiling strongly affect the maximum percent neutralization (Figure 4A), suggesting that the differential selection measures captured both neutralization-curve phenotypes.

**Figure 4.**
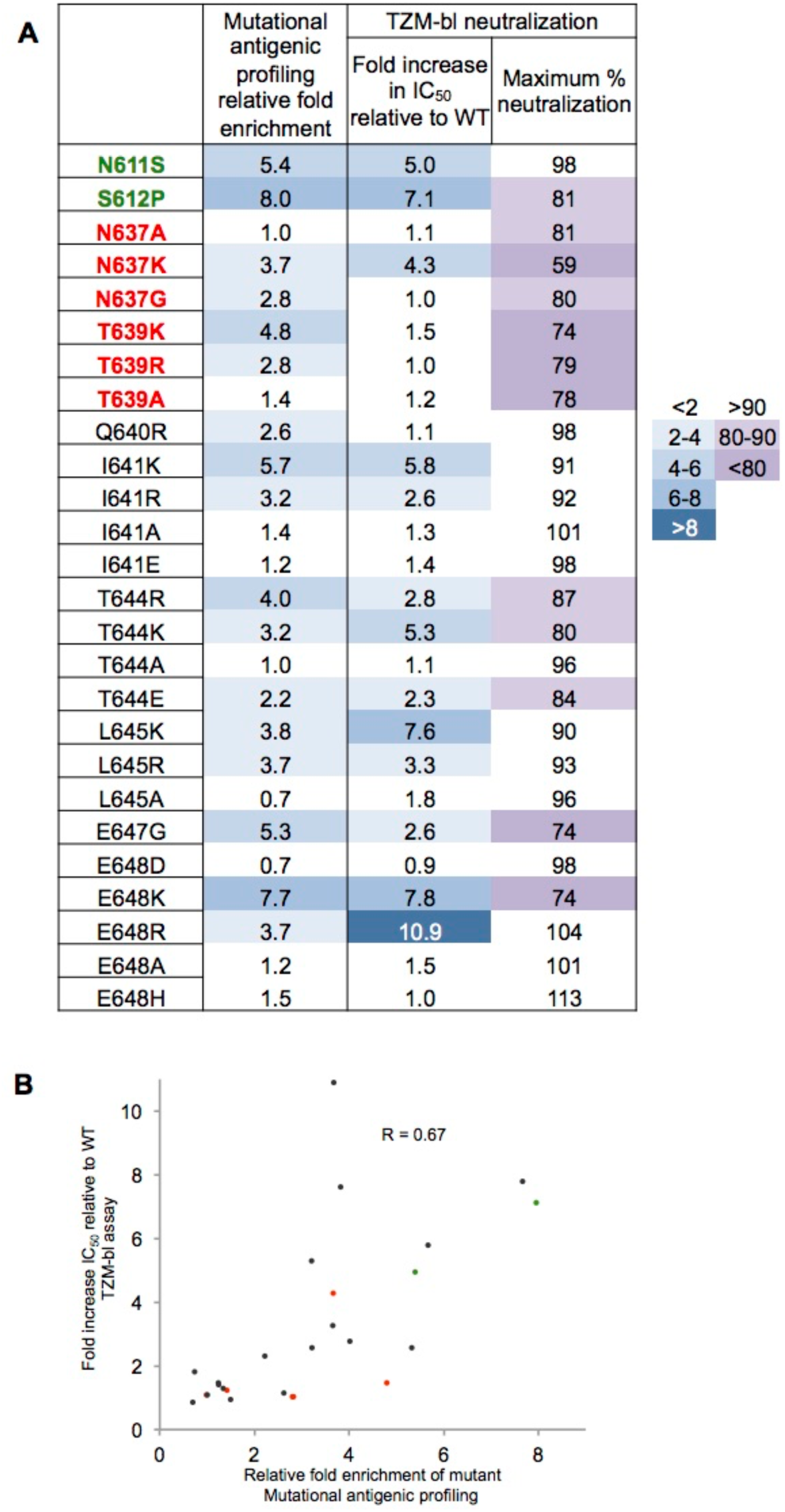
Comparison of effects of mutations in mutational antigenic profiling vs. TZM-bl neutralization assays. **(A)** Comparisons between fold-enrichment in mutational antigenic profiling of each validated mutation (Figure 3), and the fold increase in IC_50_ value relative to wildtype and maximum percent neutralization estimated from replicate neutralization assays. **(B)** For each individually tested mutations, the Pearson correlation between the fold-enrichment in mutational antigenic profiling and the neutralization assay fold-increase in IC_50_ relative to wildtype. Mutations that disrupt the 611 and 637 glycan are shown in green and red respectively.

### Comparison of our escape profiles to the original gene-wide functional mapping of PGT151

The original residue-level mapping of the PGT151 epitope was based on neutralization of two large panels of pseudovirus mutants (Falkowska et al., 2014). This work provided a large dataset against which we can benchmark our results and contrast our high-throughput measurements with results from traditional approaches. Figure 5A shows the differential selection that PGT151 exerts on every mutation at every site in Env with data from the original mapping of PGT151 shown in color bars below. In a single massively parallel experiment, we have recapitulated these original results (identifying sites 611,637, and 647), identified sites mapped in subsequent studies (sites 512 and 514), and revealed new sites of escape in the HR2 domain. In contrast, examining mutations one-by-one in neutralization assays is exceedingly low throughput - for instance, even with herculean effort (Falkowska et al., 2014), traditional approaches have only managed to test mutations at a fraction of the sites in Env (Figure 5B). The discrepancy in completeness is even more apparent when considering the fact that testing one amino-acid mutation at a site does not reveal the effects of other mutations at the same site. Whereas our study examined all 12,559 amino-acid mutations, prior work has managed to examine only a small fraction of these mutations (Figure 5B).

**Figure 5.**
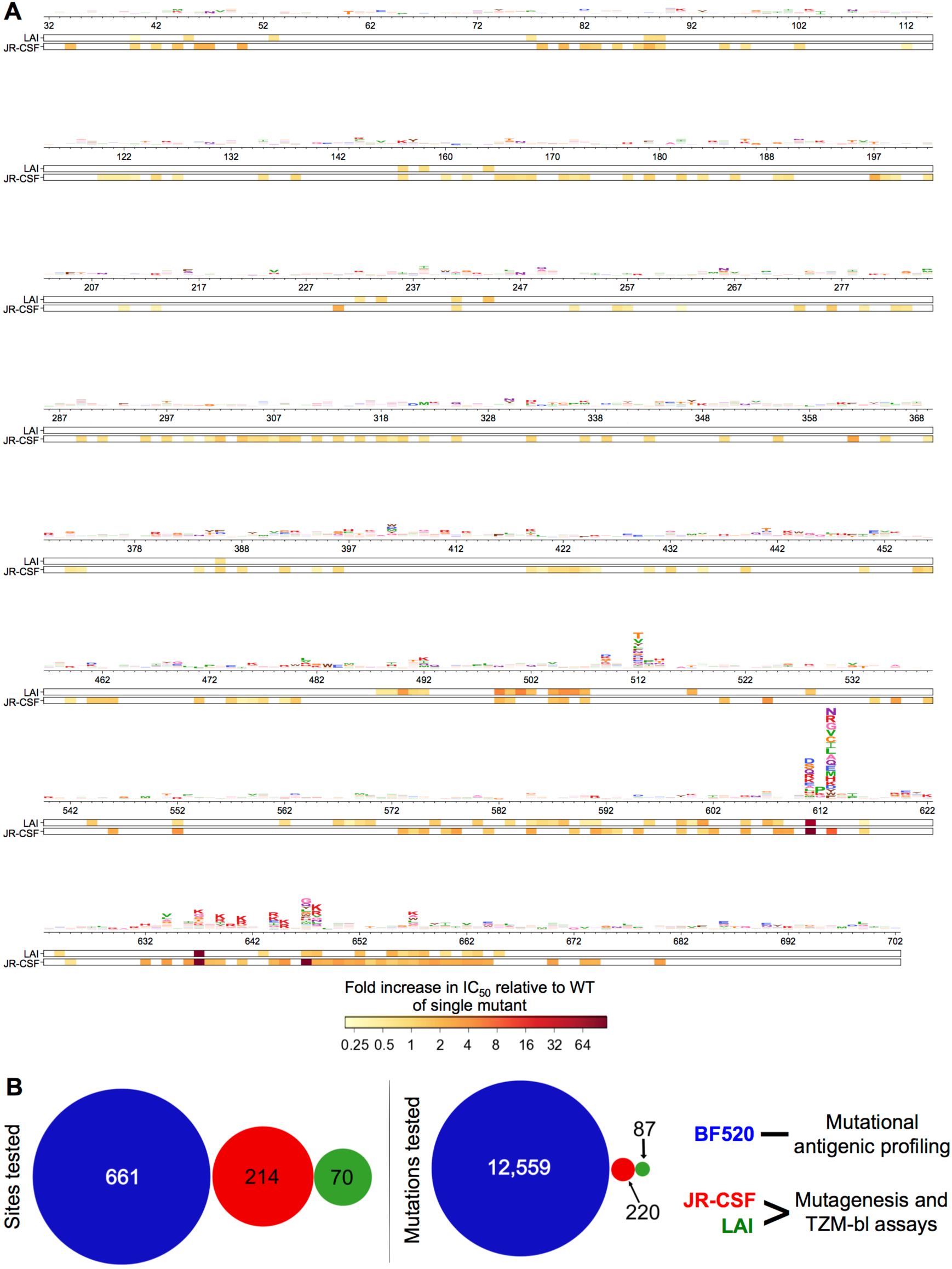
Complete mapping of escape from PGT151. **(A)** Differential selection for the entire mutagenized portion of Env. For systematic comparison, we plotted data from the original single residue-level mapping utilizing large panels of predominantly alanine scanning mutants of LAI and JR-CSF (Falkowska et al., 2014); sites where mutations were tested are colored according to the fold increase in IC_50_ relative to WT as in Figure 3. **(B)** The number of sites or mutations tested for ability to replicate and escape PGT151 neutralization in the BF520 strain using mutational antigenic profiling compared with the number of sites and mutations tested via traditional mutagenesis in the original mapping of PGT151.

### Structural insights into biochemical basis of escape from PGT151

We can interpret the differential selection measured by mutational antigenic profiling in the context of known structural information about the interface between Env and PGT151. Figure 6A maps the maximum mutation differential selection at each site onto the cryo-EM model of JR-FL Env trimer bound by PGT151 Fabs (Lee et al., 2016). The strongest differential selection maps to the structurally defined PGT151 epitope. The positively charged third complementary determining region of the heavy chain (CDRH3) of PGT151 extends into the negatively charged inter-protomer cavity at the interface between gp120 and the HR2 domain of gp41 (FIgure S5). There is differential selection predominantly for mutations to positively charged amino acids on the CDRH3-proximal side of the HR2 alpha helix (Figure 6, Figure S5), suggesting that escape in this portion of the epitope is mediated by the introduction of charge-charge repulsions.

**Figure 6.**
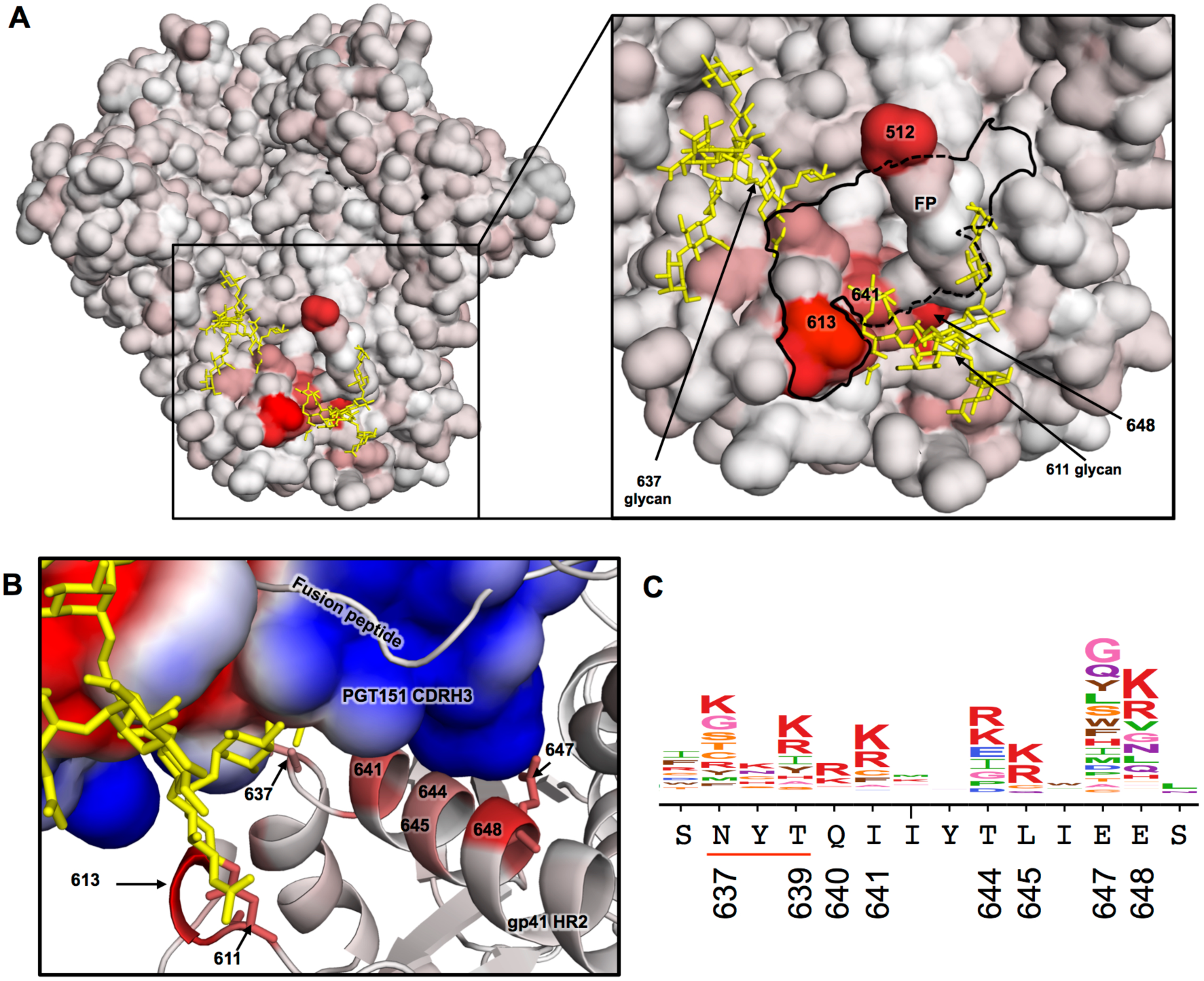
Mutational antigenic profiling combined with structural analysis suggests escape via the introduction of charge repulsions. **(A)** The JR-FL EnvΔCT trimer cryo-EM model (Lee et al., 2016) with its surface colored according to the maximum mutation differential selection at each site. The 611 and 637 glycans are shown in yellow. The inset outlines the PGT151 footprint in black, defined as residues that come within 4 angstroms of the bound PGT151 Fab, with the 611 glycosylation motif residues included for clarity. The bound PGT151 Fab is not shown, and the fusion peptide (FP), which is sequestered into a hydrophobic pocket of bound PGT 151, protrudes from the epitope in this view. **(B)** A side view of the PGT 151 CDRH3 – Env interface. Here, the ribbon representation of Env is colored according to the maximum mutation differential selection at each site, while the surface of the bound PGT151 Fab is colored according to the Poisson-Boltzmann electrostatic surface potential (red to blue; negative to positive). The negatively charged side chain of site E647 is shown in stick representation. See Figure S5 for an alternate view of this interface. **(C)** Logoplot showing the differential selection of the HR2 domain shown in **B** for reference.

The importance of charge-charge repulsions may extend to the glycosylated residue 637 (Figure S5C). While any mutation that eliminates the 637 glycan decreases the maximum percent neutralization, the introduction of a positively charged lysine results in a large reduction in both maximum percent neutralization and increase in IC_50_ (Figure 3C, 5A). Similarly, mutating site 637 to lysine has a larger effect than mutating it to alanine in multiple other strains (Falkowska et al., 2014).

Indeed, it may be possible to conceptually distinguish between sites where almost any mutation mediates escape by *eliminating* key antibody-Env interactions, versus sites where only a few mutations mediate escape by *introducing* new steric or electrostatic clashes. For instance, site 647 appears to fall into the first category, with both our mutational antigenic profiling of BF520 (Figure 6C) and prior work with JR-CSF (Falkowska et al., 2014) suggesting that many mutations mediate escape. Indeed, the cryo-EM model suggests that the E at site 647 may interact with positively charged portions of PGT151’s HCRD3 arm (Figure 6B, Figure S5C). In contrast, most of our newly identified sites of escape mutations, selection is for mutations to positively charged amino acids that are likely to clash with the positively charged antibody paratope (Figure 6B, Figure S5C). Such sites are especially difficult to identify by classical approaches such as alanine scanning, since the ability to escape neutralization is strongly dependent on which amino acid is introduced.

While the spatial clustering and biochemical evaluation of escape mutants suggests that they directly affect the interface between PGT151 and Env, it is possible that mutations indirectly affect neutralization by altering Env’s conformation. To examine this possibility, we tested some of the most strongly selected PGT151 escape mutants with other bnAbs targeting the CD4 binding site and trimer apex. None of these mutations increased sensitivity to these other bnAbs targeting distal epitopes (Table S1). This fact strongly supports the idea that most PGT151 escape mutants that we have mapped directly disrupt the antibody-Env interface.

## Discussion

We have developed a massively parallel approach to interrogate the neutralization phenotype of all functionally tolerated single amino-acid mutations to Env. We have used this approach to profile escape from PGT151, which targets a complex quaternary epitope. Our results recapitulate the escape mutations identified by previous low-throughput studies, reveal additional escape mutations not identified by these studies, and provide insight into the biochemical basis of escape.

Fine-resolution bnAb epitope mapping has previously been a cumbersome endeavor. Crystal structures of bnAb-Env complexes are often considered the gold standard, but can be difficult to obtain. In addition, the static structures do not reveal pathways of viral escape. Simply observing the evolution of HIV in the presence of a bnAb can identify specific escape mutations (Bar et al., 2016; Barouch et al., 2013; Diskin et al., 2013; Klein et al., 2012; Lynch et al., 2015a, 2015b; Shingai et al., 2013; Trkola et al., 2005), but the stochasticity of evolution means that each study only identifies one of potentially many pathways of escape. Functional residue-level mapping traditionally relies upon generating and testing Env mutants one-by-one, a resource-intensive approach that can only be applied to a fraction of Env mutations. For instance, such studies mostly restrict themselves to alanine mutations at surface sites, making them biased towards certain types of escape mutations. Therefore, despite the fact that PGT151 has been a subject of multiple studies using diverse techniques (Blattner et al., 2014; Falkowska et al., 2014; Van Gils et al., 2016; Kong et al., 2016; Lee et al., 2016; Wibmer et al., 2017), our mutational antigenic profiling has yielded a far more comprehensive and unbiased map of viral escape than all this previous work.

This complete map offers a nuanced understanding of biochemical mechanisms of escape via evaluation of the physicochemical properties of the particular amino-acid mutations that enable escape at each site. For instance, our data suggest that introduction of electrostatic clashes between the CDRH3 of PGT151 and the HR2 domain of gp41 is a previously unappreciated mechanism of escape. The specificity of escape at these sites contrasts with the many diverse amino-acid mutations that were enriched at residue 647. This broad escape profile suggests that elimination of a key Env-PGT151 interaction underlies escape at site 647. The systematic quantification of the effect of each amino-acid mutation at each site has the potential to augment structural studies by providing insight into the energetics of antibody-epitope interactions.

The ability to comprehensively map viral escape also has important application for bnAb-based immunotherapies and vaccines. Quantifying how epitope features contribute to neutralization could aid in engineering broader and more potent antibodies (Diskin et al., 2011). Such information could also inform the design of immunogens that elicit responses that thwart common pathways of viral escape. Completely defining viral determinants of escape will also aid in evaluating the efficacy and failure of bnAb immunotherapeutics in humans, just as determining drug resistance mutations has aided the development of antiviral therapy and prophylaxis (Lehman et al., 2015; Tural et al., 2002). Similar to algorithms that leverage large datasets of drug-resistance mutations to predict antiviral resistance based on viral genotype (Vercauteren and Vandamme, 2006), comprehensive mutation-level escape profiles could inform similar sequence-based scoring metrics for bnAb resistance.

As with all epitope-mapping approaches, mutational antigenic profiling has limitations. We are evaluating the effects of single amino-acid mutations to a single viral strain. However, strain-specific differences in bnAb sensitivity have been described for PGT151 (Blattner et al., 2014; Falkowska et al., 2014), and could explain why we did not observe escape at sites (533, 537, and 540) that have been shown to affect the sensitivity of JR-CSF to PGT151 (Van Gils et al., 2016). Future mutational antigenic profiling of bnAb escape in multiple different Envs could determine the prevalence and mechanisms of such strain-specific differences. Additionally, mutational antigenic profiling of different antibodies could identify mutations that globally alter neutralization sensitivity. While we did not observe evidence of allosteric or global escape mutations in our work with PGT151, unbiased examination of all mutations should reveal such mutations when they exist.

Few protein-protein interactions have been as heavily studied as those between bnAbs and Env. Indeed, these interactions provide the motivation for many current HIV treatment and prevention efforts. We have provided the first complete map of the viral determinants of neutralization and escape at one of these bnAb-Env interfaces. The mutational antigenic profiling approach that we have used to obtain this map is high-throughput and quantitative. We anticipate that this approach can be extended to define all possible HIV escape mutations from other bnAbs and possibly polyclonal serum. The resulting maps could be valuable for informing the design of immunogens and the development and evaluation of bnAb immunotherapies.

## Experimental Procedures

### Data and source code

An iPython notebook containing our analysis is included as File S1. The software used to align sequence reads and compute differential selection is available at https://github.com/jbloomlab/dms_tools (Bloom, 2015). The deep sequencing data are available on the Sequence Read Archive under accession numbers SRX2548567- SRX2548579, and the mutation differential selection values are provided as File S2.

### Generation of mutant DNA libraries

Codon mutant libraries were created in the context of BF520 *env* introduced into the full-length proviral clone Q23 (Poss and Overbaugh, 1999). Q23 is a subtype A transmitted/founder virus that is the basis for a well characterized system of generating chimeric full length proviral clones as well as pseudoviruses encoding heterologous *env* genes (Provine et al., 2009).

We first introduced codon mutations into *env* in three independent replicates using a slightly modified version of the PCR mutagenesis technique as previously described (Bloom, 2014). To overcome potential biases in the frequency of introduced mutations, we used mutagenesis primers of equal melting temperature rather than equal length. These primers are provided as File S4, and the script to generate these is available at https://github.com/jbloomlab/CodonTilingPrimers. We then cloned 4×10^5^ – 1×10^6^ unique variants of for each mutant *env* library into a high efficiency-cloning vector (Q23.BsmBI.∆Env, File S5) that utilizes BsmBI restriction sites to clone *env* into the Q23 backbone. T4 DNA ligation products were transformed into ElectroMAX DH10B competent cells (Invitrogen; 12033-015). The next day, plated colonies were scraped, grown in LB plus ampicillin for 4 hours, and maxiprepped. The number of unique variants per library was determined by counting colonies on plates containing a dilution of the high efficiency transformation.

### Generation of mutant virus libraries

To generate mutant virus libraries, 36 ug total of each mutant DNA library was transfected using Fugene-6 into 18 wells of 6-well plates, that had been plated with 4×10^5^ 2 93T cells per well the previous day. Two days post transfection, supernatants were collected, filtered through a 0.2 uM filter, and DNAse (Roche; 4716728001) treated to eliminate leftover transfection plasmid as previously described (Haddox et al., 2016). This transfection supernatant was then titered on TZM-bl cells by adding serial dilutions of the supernatant to 20,000 cells in the presence of 10 μg/mL DEAE-dextran in 600 μL total volume. After 48 hours, cells were fixed and stained for beta-galactosidase, and infected cell foci were counted.

To establish a genotype-phenotype link and select for functional variants, we passaged each mutant library for 4 days in SupT 1.CCR5 cells at a low MOI (0.01 TZM-bl infectious units/cell). This initial passage in in SupT1.CCR5 cells was performed similarly to (Haddox et al., 2016). We infected cells in with 3×10^6^ (replicate 1) or 4×10^6^ (replicates 2 and 3) infectious units of transfection supernatant in T-225 flasks, each with 100 mL total volume of R10 (RPMI [GE Healthcare Life Sciences; SH30255.01], supplemented with 10% FBS, 1% 200 mM L-glutamine, and 1% of a solution of 10,000 units/mL penicillin and 10,000 μg/mL streptomycin) in the presence of 10 ug/mL DEAE-dextran with cells starting at a concentration of 1×10^6^ cells/mL. On day 1, we replaced the media with fresh R10 containing 10 ug/mL DEAE dextran, and on day 2, we doubled the total volume, splitting each flask into two. On day 4, we pooled all the flasks, spun down cells, and filtered the media through a 0.2 uM filter. We then concentrated this virus ~33 fold via ultracentrifugation for 1 hour at 4°C at 23,000 RPM over a 20% sucrose cushion using an SW 28 rotor (Beckman Coulter; 342207) and resuspended in R10. In parallel for each replicate, we passaged 5×10^5^ infectious units of wildtype virus under the same conditions as a control. These passaged viruses were titered on TZM-bl cells.

### PGT151 selection of mutant virus libraries

We incubated each mutant virus library with PGT151 and infected SupT1.CCR5 cells again. Each library was also passaged without PGT151 treatment to serve as a replicate-specific control to calculate differential selection. For each condition, 10^6^ TZM-bl infectious units of each library was incubated +/-1 ug/mL of PGT151 at 37°C for 1 hour, then infected into 10^6^ (not PGT151 treated) or 2×10^5^ (PGT151 treated) SupT1.CCR5 cells in the presence of 100 ug/mL DEAE dextran. Three hours post infection, cells were spun down and resuspended in fresh R10, containing no DEAE-dextran. At 12 hours post infection, cells were spun down, washed with PBS, and then subjected to a mini-prep to harvest non-integrated viral cDNA. We also infected non-neutralized wildtype virus in parallel for each library.

### Deep sequencing

To determine the frequency of each mutation in the antibody-selected and nonselected conditions, we utilized a barcoded subamplicon Illumina deep sequencing approach as previously described (Doud and Bloom, 2016; Haddox et al., 2016). This approach uses unique molecular identifiers to distinguish true mutation from sequencing errors. It reduced the error rate when sequencing wildtype proviral plasmid to 1.5×10^-4^ mutations per codon (Figure S2). Data were analyzed using dms_tools (Bloom, 2015) version 1.1.20 as described in the computational pipeline provided in File S1. This pipeline details sequencing depth and the number of unique molecules sequenced for each amplicon for each sample (roughly 1×10^5^ - 3×10^5^ unique error-corrected molecules per amplicon for PGT151 selected samples). Primers used during the sequencing library preparation are provided in File S6.

### Computation of differential selection

We calculated differential selection values as described in (Doud et al., 2017). Briefly, the enrichment (*E_r_*_,*x*_) of each amino acid ! at site *r* relative to wildtype is calculated as shown in *equation 1,* where 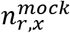 is the number of counts of *x* at site *r* in the mock treated sample. Similarly, 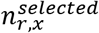 is the number of counts of *x* at site *r* in the PGT151 selected sample. *wt*(*r*) is the wildtype character at *r.*

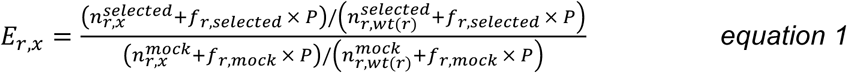

To account for statistical noise associated with low counts, a pseudocount of *P* =20 was added to each count. The *f_r,selected_* (*equation 2*) and *f_r_,_mock_* (*equation 3*) variables scale the pseudocount to account for different sequencing depth of the *mock* and *selected* libraries at site *r.*

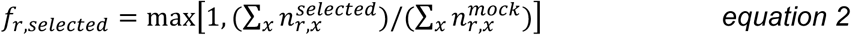

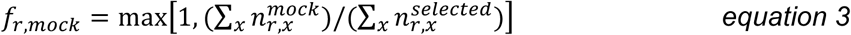

To account for errors in viral replication and sequencing, counts in *equation 1* were adjusted by the rates of mutation to *x* at site *r* in mock selected wildtype virus libraries passaged in parallel for each replicate. We define 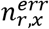 as the number of counts of *x* at site *r* in the matched wildtype virus control and *ϵ_r,x_* as shown in *equation 4,* such that *ϵ_r,x_* is the rate of errors to *x* at site *r* when *x* ≠ *wt*(*r*), and *ϵ_r,x_* is one minus the rate of errors away from wildtype at site *r* when *x = wt*(*r*) as defined in *equation 4.*

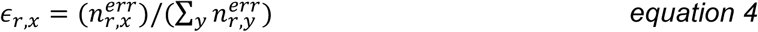

The observed counts in *equation 1* are then adjusted to the error-corrected counts *n̂_r_,_x_* as described in *equation 5.*

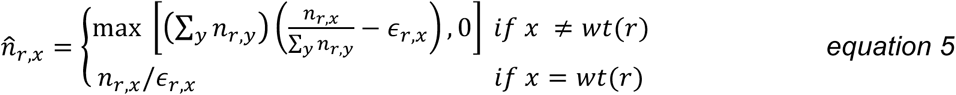

Lastly, differential selection values *s_r,x_* for *x* at *r* in the *selected* versus *mock* condition is quantified as *s_r,x_ =* log_2_*E_r,x_,* and visualized on logoplots rendered by dms_tools via weblogo (Crooks et al., 2004). Throughout this manuscript, we focused only on positively enriched mutations.

### Supplemental Experimental Procedures

Descriptions of the *env* sequence numbering, PGT151 production, TZM-bl neutralization assays, and structural analyses are provided in the Supplemental Experimental Procedures.

## Author Contributions

Conceptualization: ASD, JO, JDB

Methodology: ASD, HKH, JDB

Validation: ASD

Investigation: ASD

Software: ASD, JDB

Supervision: JO, JDB

Writing – Original Draft: ASD

Writing – Review & Editing: ASD, HKH, JO, JDB

Funding Acquisition: JO, JDB

## Acknowledgments

We thank James Hoxie for the SupT 1 .CCR5 cells, Michael Doud and Michael Emerman for helpful discussions, and Mark Pankau for assistance with ddPCR. We thank the Fred Hutch Genomics core for performing the Illumina sequencing. ASD was supported by a NSF Graduate Research Fellowship (DGE-1256082). HKH was supported by a Cell and Molecular Biology Training Grant (T32GM007270) and a Molecular Biophysics Training Grant (T32GM0008268). This work was supported by NIH grants R01GM102198 and R01AI127893 to JDB and DA039543 and R01AI120961 to JO. The research of JDB was also supported in part by a Faculty Scholar Grant from the Howard Hughes Medical Institute and the Simons Foundation.

We have no conflicts of interest.

## Supplemental Experimental Procedures

### Env Sequence numbering

Unless otherwise stated, Env residues are numbered according to the HXB2 reference strain numbering system (Korber et al., 1998). The corresponding sequence numbering based on aligned BF520.W14M.C2 (GenBank accession number KX168094.1) and HXB2 *env* for the mutagenized portion of the gene is provided as File S3.

### PGT151 production

PGT151 heavy (GenBank: KJ700282.1) and light (GenBank: KJ700290.1) chains were codon optimized, cloned into Igγ1 and Igκ expression vectors, and expressed in 293F cells using the FreeStyle MAX system (Invitrogen). IgG was purified using Protein G resin (Pierce; 20399) as described by Simonich et al., 2016.

### ddPCR protocol

To quantify the remaining infectivity of each mutant virus library after PGT151 selection and infection into SupT1.CCR5 cells, we used droplet-digital PCR. The number of viral genomes present in each harvested sample of viral cDNA was quantified using a *pol* PCR (Benki et al., 2006) adapted for digital droplet detection (Strain et al., 2013). The percent escape shown in Figure 2 was calculated using the number of genomes present in each selected library relative to its non-neutralized control.

### TZM-bl neutralization assays

To validate our escape profiles, we generated pseudoviruses bearing individual point mutants of both enriched and non-enriched mutations. We then performed TZM-bl neutralization assays as previously described (Cortez et al., 2015) in the presence of 10 ug/mL DEAE-dextran. Neutralization assays were performed in duplicate, and WT neutralization curves were run on each plate to reduce noise (Figure S4). Neutralization curves were fit using 3-parameter nonlinear regression of the % neutralization values across the dilution series, with the bottom plateau constrained to 0 (Figure S4). To calculate the IC_50_ relative to wildtype, curves were solved for y = 50% neutralization. The fitted top plateau, averaged across replicates, is the maximum percent neutralization. Fold change in IC_50_ relative to WT was calculated within a single experiment and then averaged across replicates.

### Structural analyses

All structural analyses are based on the cryo-EM model of JR-FL EnvΔCT trimer bound by two PGT151 Fabs ((Lee et al., 2016); PDB ID: 5FUU). For all figures, we focused on PGT151-Env interface 2 (Lee et al., 2016). Figures were generated using Pymol, and the Poisson-Boltzman electrostatic surface potential was calculated using the APBS plugin (Baker et al., 2001) with the protein dielectric constant of 20 for the trimer or PGT151 Fab (trimer bound conformation) in isolation.

---

**Figure S1.**
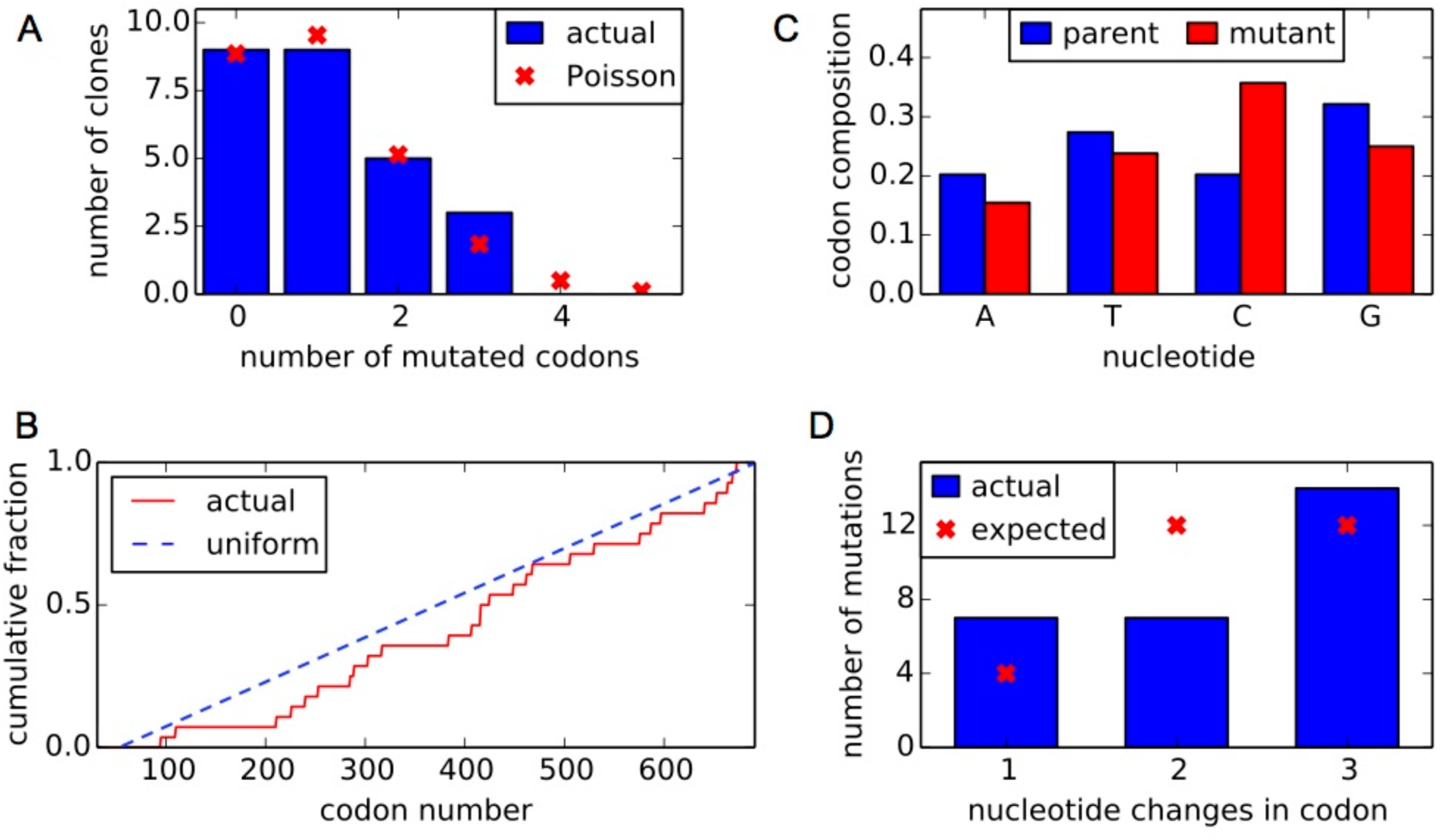
Mutant DNA library statistics based on Sanger sequencing data from 26 individual clones. Related to Figure 1 and Experimental Procedures. Clones were obtained from the three replicate mutant libraries of the proviral plasmid (Fig 1C). **(A)** The distribution of codon mutations per clone is shown, comparing the actual number with the expect number based on a Poisson distribution. **(B)** The cumulative distribution of mutations across the length of the mutagenized portion of the gene compared to the expected uniform distribution. In this panel, codons are numbered sequentially along the Env sequence beginning with 1 at the N-terminal methionine. **(C)** The nucleotide composition of parent and mutant codons for each mutation. The introduced mutations had relatively uniform nucleotide composition. **(D)** The average number of nucleotide changes of each codon mutation compared to the expected if each codon is equally likely to be mutated to every other codon. Our codon mutagenesis introduced a mix of 1-nucleotide (e.g., GCA -> GtA), 2-nucleotide (e.g., GCA -> Gtc), and 3-nucleotide (e.g., GCA -> atc) codon mutations.

**Figure S2.**
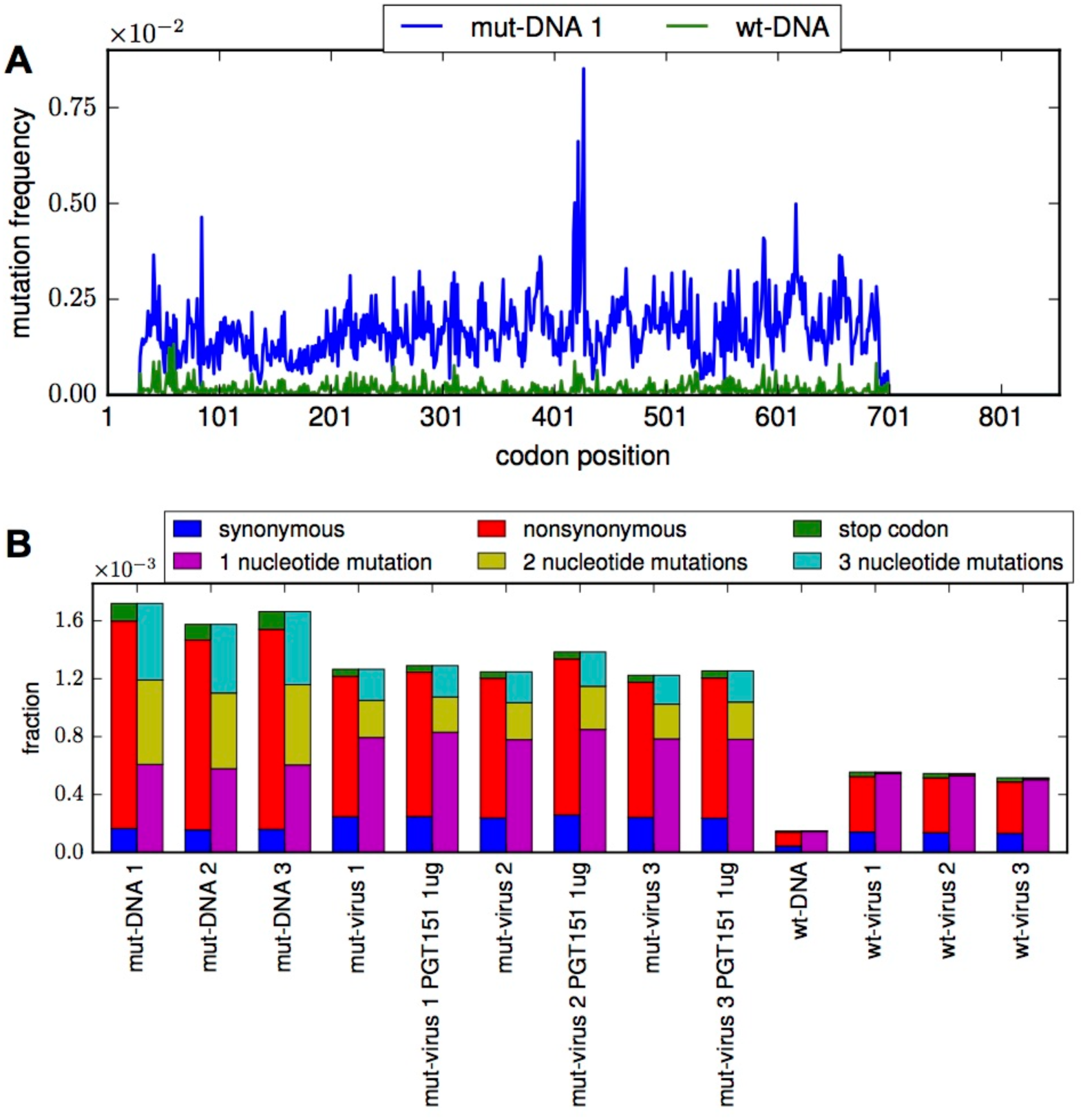
Mutation frequencies of wildtype and mutant DNA and virus libraries. Related to Figure 1 and Experimental Procedures. **(A)** The codon mutation frequency across the gene is, on average, >10 times higher in the mutant DNA libraries (*mut-DNA*) than the error frequency observed when sequencing wildtype proviral plasmid (*wt-DNA*). In this panel, codons are numbered sequentially along the Env sequence beginning with 1 at the N-terminal methionine. **(B)** The frequency of mutation types in the wildtype and mutant DNA and virus libraries. Compared to wildtype proviral plasmid (*wt-DNA*), there is an increase in single nucleotide mutations in the wildtype passaged virus libraries (*wt-virus*), presumably from errors during viral replication. There is also evidence of purging of deleterious mutations (e.g. stop codons) from the mutant plasmid libraries (*mut-DNA*) relative to the mutant viruses (*mut-virus*) grown from these plasmids, both in the presence and absence of PGT151 antibody.

**Figure S3.**
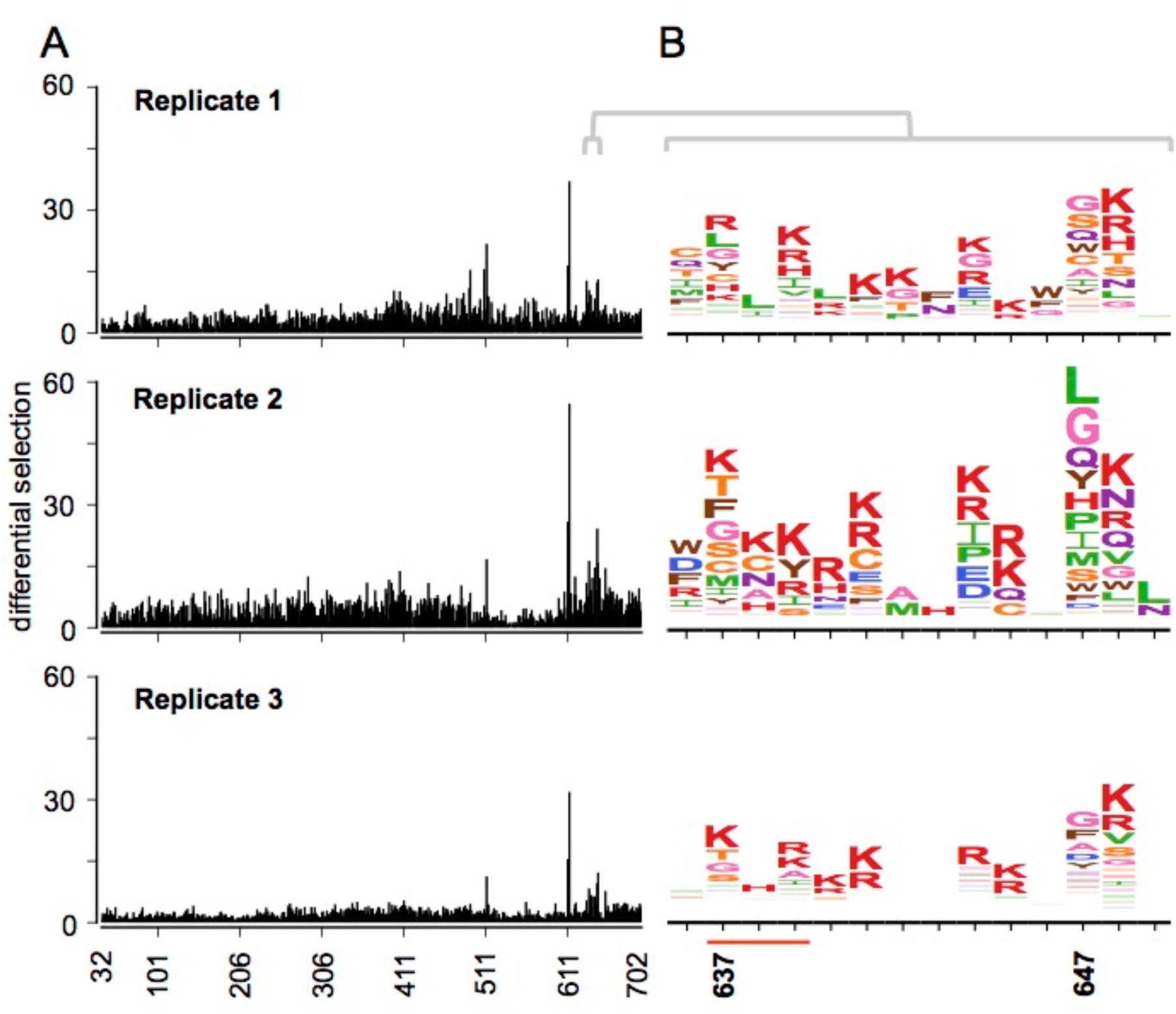
Reproducibility of mutational antigenic profiling. Related to Figure 2. **(A)** The site differential selection is plotted as in Figure 2C, but separately for each replicate. **(B)** The escape profile of the 637 glycan motif (underlined in red) and HR2 domain is shown individually for each replicate. All logoplots are plotted on the same scale.

**Figure S4.**
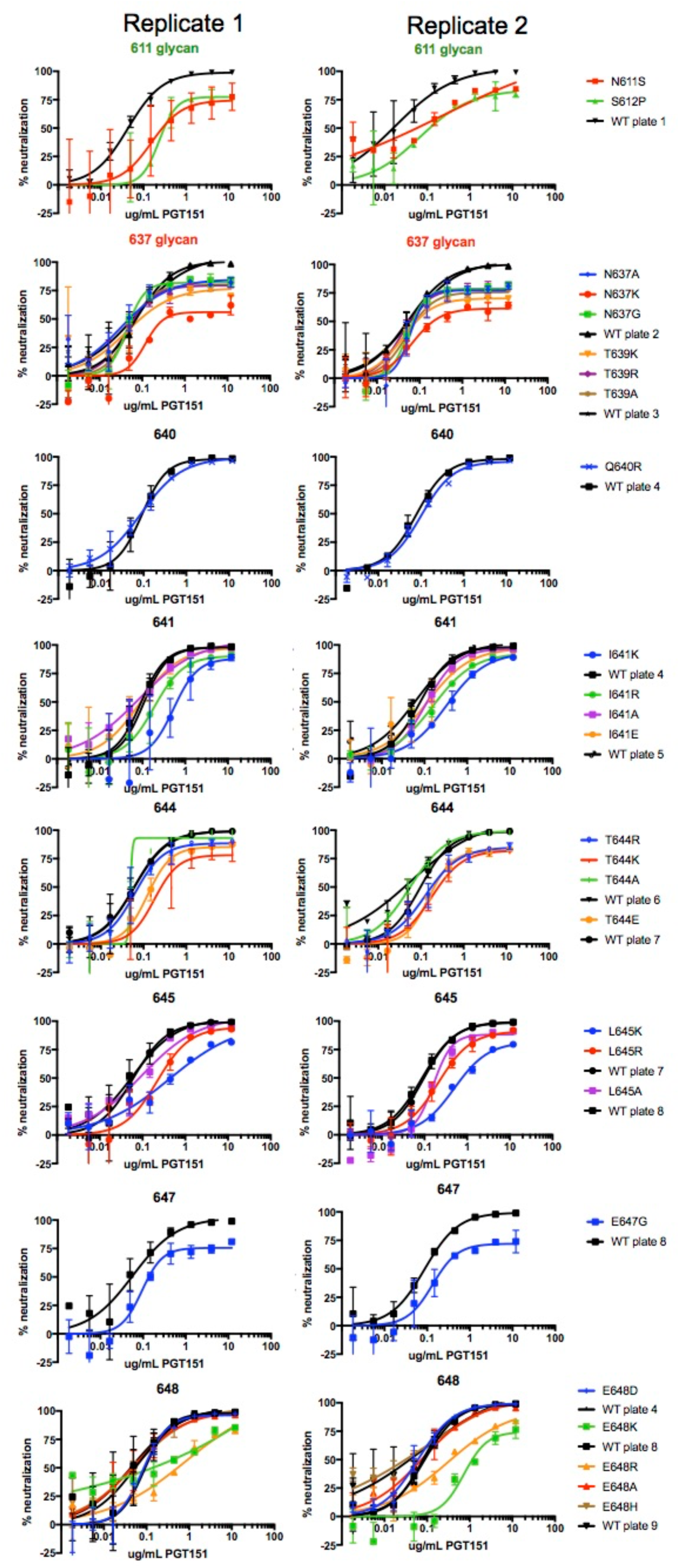
Replicate TZM-bl neutralization curves. Related to Figure 3 and 4. Regression curves used to estimate IC_50_ and maximum percent neutralization are plotted for each replicate. To reduce noise, the fold change in IC_50_ relative to wildtype was compared to a wildtype control run on the same plate as the mutant. When multiple mutants at a site were spread across different plates, we plotted both wildtype curves on that site’s neutralization curve plot. The wildtype curve matched to each mutant is the first wildtype curve directly beneath the mutant in the legend.

**Figure S5.**
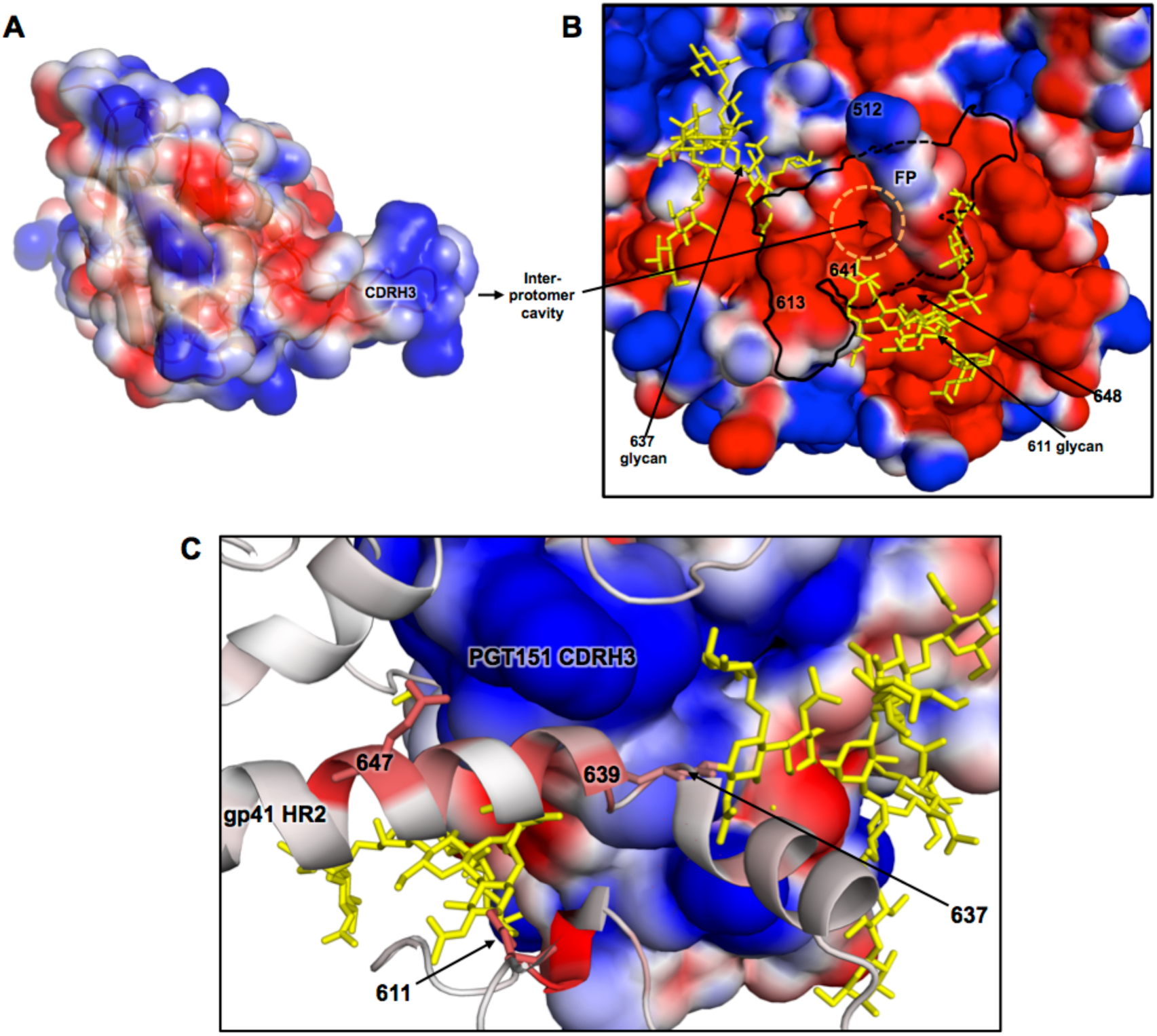
Electrostatic complementarity between PGT151 and the Env. Related to Figure 6. **(A)** The PGT151 Fab (in its trimer bound conformation) is colored according to the Poisson-Boltzmann electrostatic surface potential (red to blue; negative to positive). **(B)** The surface representation of Env as in the inset of Figure 6A, but with the surface colored according to the electrostatic surface potential. The negatively charged inter-protomer cavity that the positively charged CDRH3 arm extends into is circled in orange. **(C)** An internal view of the PGT151 CDRH3 – Env interaction, as shown in Figure 6B. Again, the ribbon representation of Env is colored according to the maximum mutation differential selection at each site, while the surface of the bound PGT151 Fab is colored according to the electrostatic surface potential.

**Supplemental Table 1.**
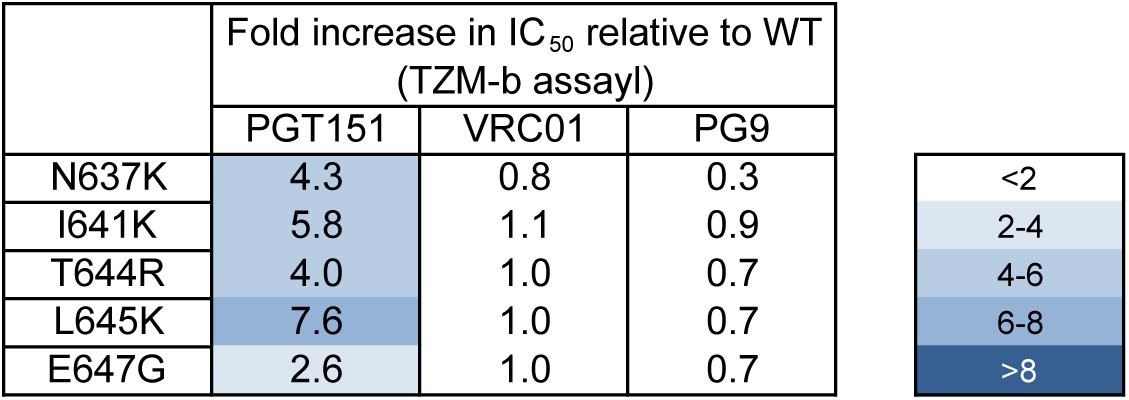
Sensitivity of BF520 PGT151 escape mutants to bnAbs targeting other epitopes. Related to Figure 6. The fold increase in IC_50_ relative to wildtype for each mutant is shown for PGT151, VRC01 (targets the CD4 binding site) and PG9 (targets glycans at trimer apex). Fold increase in IC_50_ relative to WT is the averaged over two independent TZM-bl assays. Additionally, numerous other previously identified PGT151 escape mutants (also characterized in BF520 in this study) have been shown to not appreciably affect neutralization in other strains by a number of bnAbs targeting distal epitopes (Van Gils et al., 2016; Kong et al., 2016; Wibmer et al., 2017).

